# Highly Efficient Antibiofilm and Antifungal Activity of Green Propolis against *Candida* species in dentistry material

**DOI:** 10.1101/2020.01.27.920959

**Authors:** Carolina Rabelo Falcão Bezerra, Katia Regina Assunção Borges, Rita de Nazaré Silva Alves, Amanda Mara Teles, Igor Vinicius Pimentel Rodrigues, Marcos Antonio Custódio Neto da Silva, Maria do Desterro Soares Brandão Nascimento, Geusa Felipa de Barros Bezerra

## Abstract

**Background:** This study evaluated the influence of green propolis’ extract on the adhesion and biofilm formation of *Candida* species on dentistry material.

**Methods:** Phytochemical analysis of green propolis’ extract was performed by High Performance Liquid Chromatography. Adhesion was quantified in a Neubauer chamber, counting the number of yeast cells adhered to the fragments; Biofilm formation was determined by counting the number of colony forming units (CFU). The intensity of biofilm formation adhesion was classified as negative, weak, moderate, strong and very strong. Fifteen compounds were identified in green propolis extract, mainly flavonoids.

**Results:** All strains were able to adhere and form biofilm on the surface of the orthodontic materials studied. In steel and resin, the adhesion intensity of the yeast cells was weak at all incubation times, except for *C. parapsilosis* and *C. tropicalis* which at 12hs showed moderate intensity. Regarding biofilm formation (24 and 48 hours), it was observed in the steel that *C. albicans* had moderate intensity at 24 and 48 hours; *C. parapsilosis* at 24 and 48 hours had very strong intensity; *C. tropicalis* at 24 hours had strong intensity and at 48 hours very strong. While in the resin, all species at 24 and 48 hours had strong intensity, except for *C. tropicalis* which at 48 hours had very strong intensity. Green propolis extract showed antifungal activity and was able to inhibit both adhesion and biofilm formation at 2.5 μg/mL.

**Conclusions:** This study reinforces the idea that green propolis has antifungal activity and interferes with virulence factors of *Candida* species.

## BACKGROUND

In recent years the use of orthodontic materials has increased for aesthetic, surgical and biofunctional purposes. Polymers, ceramics, composites, resin, steels and their alloys are used in the manufacture of dental prostheses, screws and orthodontic appliances and when implanted in the oral cavity they are exposed to colonization and biofilm formation by microorganisms that live in the oral cavity. Alongside with the pH and saliva, these devices are targets of biofilm formation especially produced by *Candida* spp. (1).

A combination of factors contribute to *Candida* sp biofilm formation, salivary flow, low pH, poor oral hygiene and the type of orthodontic material contribute to biofilm colonization and formation (2). During colonization and biofilm formation, oral microbiota secrete enzymes and exopolysaccharides to colonize a surface, thus the biofilm constitutes as a film of organic components that are absorbed from saliva forming an extracellular polymeric matrix and thus the multicellular community (bacteria or fungus) is incorporated into the extracellular matrix (ECM) (1–3).

The formation of biofilm in orthodontic materials raises concern as, when installed, increases the risk of infection, antibiotic and antifungal resistance, becoming an infectious site and obstacle for therapies. Natural products may inhibit biofilm formation, however, antibiofilm effects depends on inhibition of extracellular matrix formation, adhesin inhibition and cell attachment and inhibition of virulence factors (3).

Propolis is a resin and a natural product with medicinal properties. The production of propolis occurs from the collection of plant structures and its mixture with wax and salivary enzymes, having the modeling function of a varnish, besides protecting and sterilizing the internal and external parts of the hive, keeping the humidity and temperature (4–6).

Brazil has at least thirteen distinct types of propolis and many bioactive compounds, such as apigenin, artepilin C, vestitol, neovestitol, among others (7). There are varieties of propolis: red, green, yellow, brown, according to the flowering period. Green propolis is usually obtained from *Baccharis dracunculifolia* as a sticky exudate from leaves, flower buttons, buds, stems and fruits (8). This substance is rich in compounds such as prenylated phenylpropanoids, triterpenoids, benzoic and chlorogenic acids.

Scientific literature reports that green propolis has antifungal and antibacterial activities against *Lasiodiplodia theobromae* (9), *Candida* spp. (10) and *Streptococcus mutans* (11), *Streptococcus acidominimus*, *Streptococcus oralis*, *Staphylococcus epidermidis*, *Veillonella parvula*, *Bifidobacterium breve*, *Bifidobacterium longum*, and *Lactobacillus acidophilus*, respectively (12). Thus, the aim of this research was to evaluate the activity of green propolis extract on the virulence factors (adhesion and biofilm) of *Candida albicans*, *C. tropicalis* and *C. parapsilosis* in dental materials (acrylic resin and steel).

## RESULTS

### Phytochemical screening

In the present study, the extract showed strong reaction for flavones, flavonoids and xanthones, the average intensity reaction for the presence of alkaloids, condensed tannins, hydrolysable tannins is showed on the table below (Table 1).

**Table 1.**
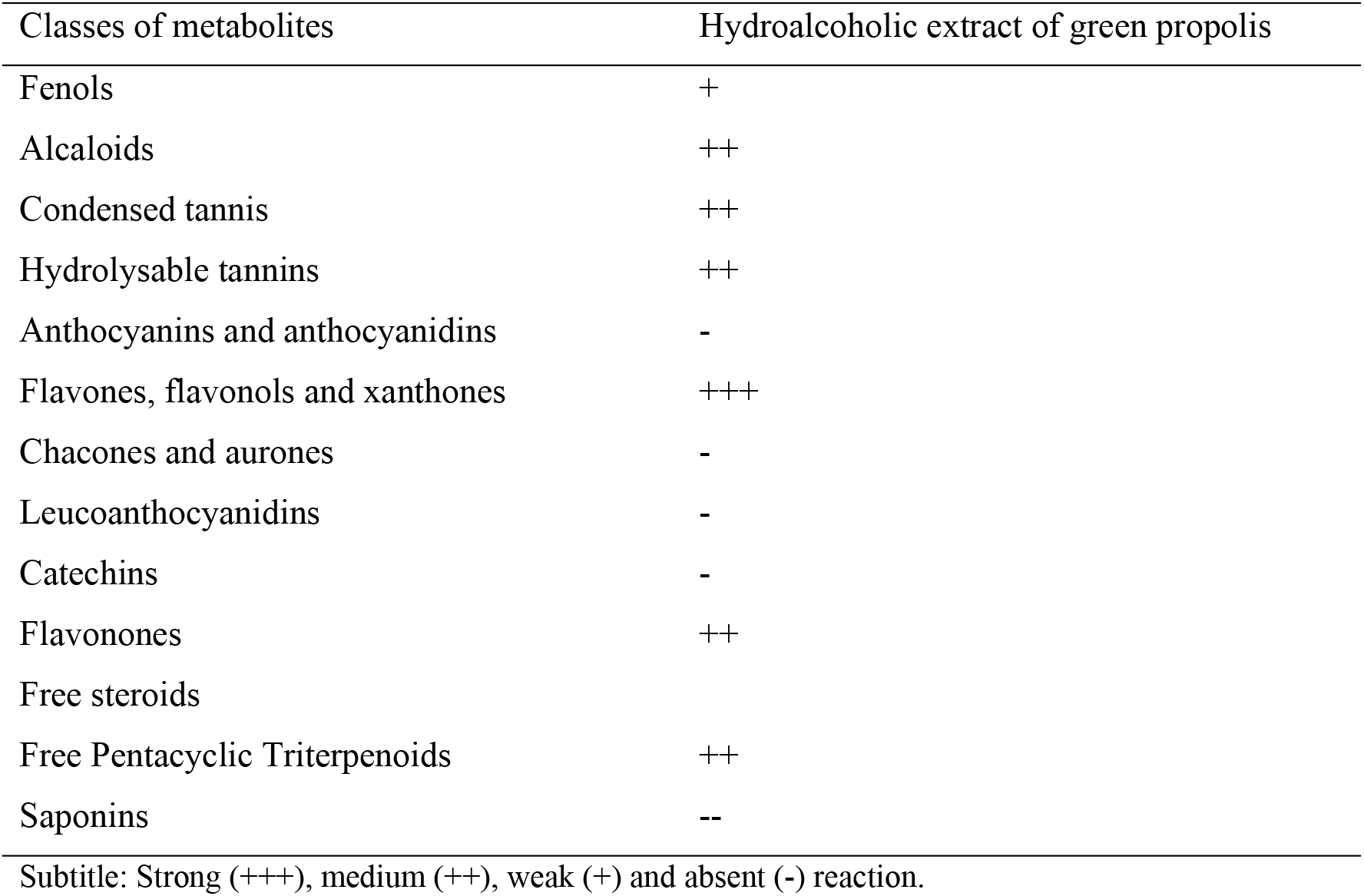
Classes of secondary metabolites identified in Green Propolis Extract.

### Chemical composition of green propolis hydroalcoholic extract by HPLC-DAD-MS

The profile of the compounds was analyzed by HPLC-DAD-MS (Figure 1). Fifteen compounds were identified in the green propolis extract (Table 2). The main compounds are flavonoids and phenolic acids.

**Figure 1.**
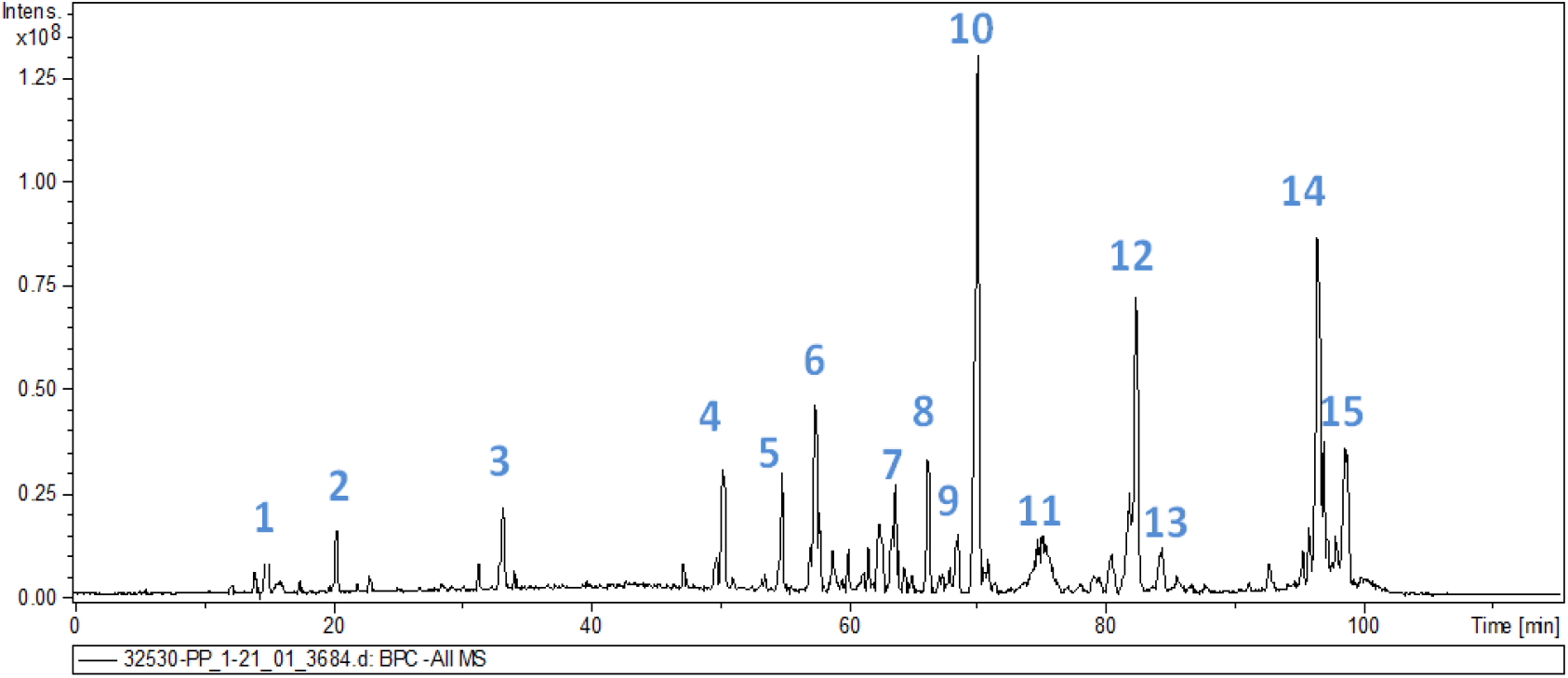
Chromatogram of the hydroalcoholic green propolist extract at a wavelength of 270 nm.

Isolated chemical compounds are described in Table 3, including retention time and observed mass. The spectra of each peak identified on the HPLC-DAD-MS are described in the supplementary article material. The chemical structures are described in Table 4, with their masses. The spectra of the fifteen compounds are described in the supplement material.

**Table 3.**
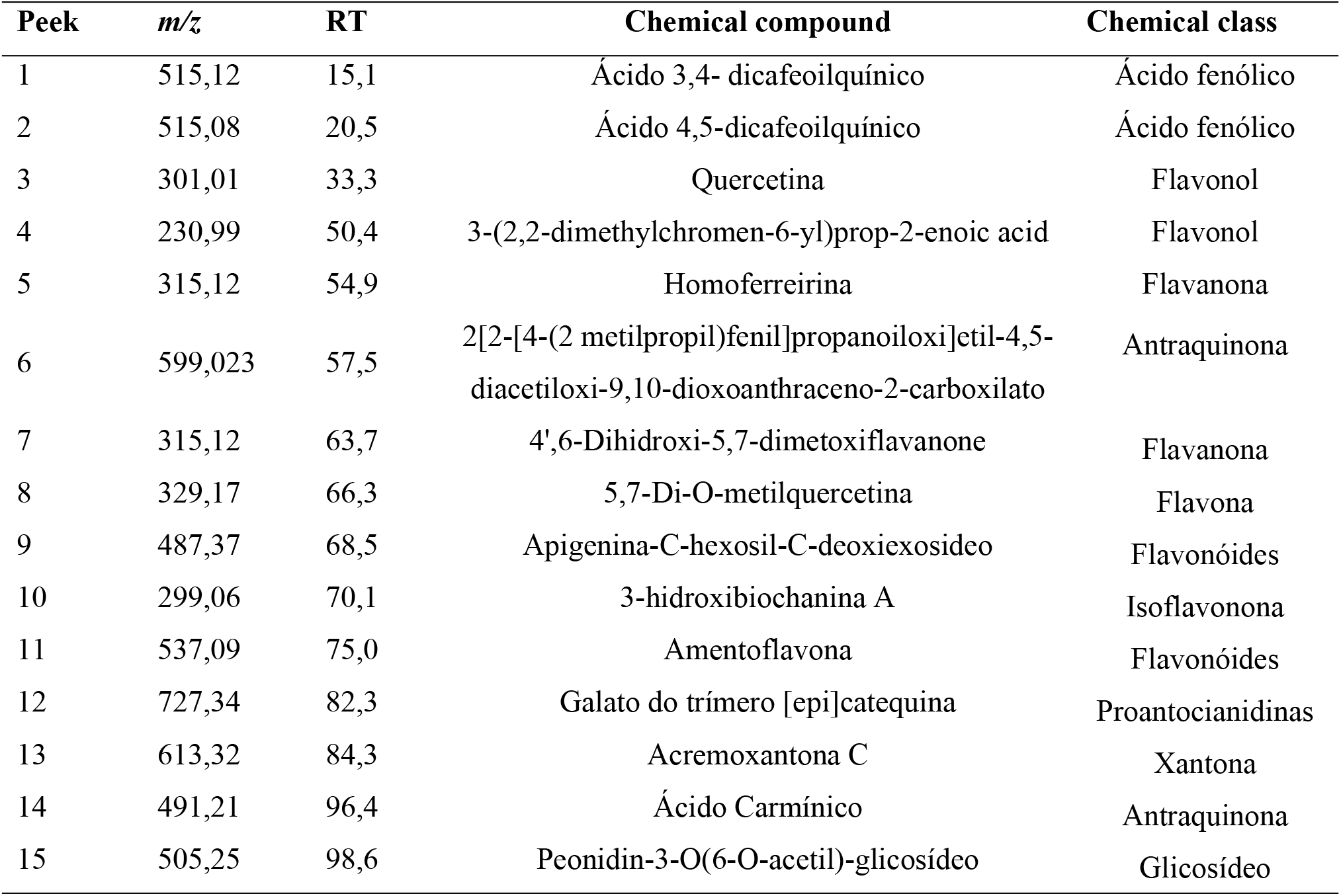
Chemical compounds, mass, retention time (RT) from green propolis by HPLC-DAD-MS

**Table 4.** Chemical compounds identification in green propolis by HPLC-DAD-MS

| Chemical compound | Structure | *m/z* |
| --- | --- | --- |
| 1 | C<sub>25</sub>H<sub>24</sub>O<sub>12</sub> | 515,12 |
| 2 | C<sub>25</sub>H<sub>24</sub>O<sub>12</sub> | 515,08 |
| 3 | C<sub>15</sub>H<sub>10</sub>O<sub>7</sub> | 301,01 |
| 4 | C<sub>14</sub>H<sub>14</sub>O<sub>3</sub> | 230,99 |
| 5 | C<sub>17</sub>H<sub>16</sub>O<sub>6</sub> | 315,12 |
| 6 | C<sub>34</sub>H<sub>32</sub>O<sub>10</sub> | 599,023 |
| 7 | C<sub>17</sub>H<sub>16</sub>O<sub>6</sub> | 315,12 |
| 8 | C<sub>16</sub>H<sub>14</sub>O<sub>7</sub> | 329,17 |
| 9 | NI | 487,37 |
| 10 | C<sub>16</sub>H<sub>12</sub>O<sub>5</sub> | 299,06 |
| 11 | C<sub>30</sub>H<sub>18</sub>O<sub>10</sub> | 537,09 |
| 12 | NI | 727,34 |
| 13 | C<sub>33</sub>H<sub>26</sub>O<sub>12</sub> | 613,32 |
| 14 | C<sub>22</sub>H<sub>20</sub>O<sub>13</sub> | 491,21 |
| 15 | C<sub>24</sub>H<sub>25</sub>O<sub>12</sub> | 505,25 |

NI= not identified

### Evaluation of antioxidant activity of green propolis extract by DPPH

The relation between the antioxidant activity (%) and the concentrations of the extract shown in the equation of the line (Y = 0.2714x + 27.966), with an R2 = 0.983 showed that the antioxidant percentage increases proportionally to the increasing concentrations of the extract, reaching 97.99% of antioxidant activity at a concentration of 275 μg / mL providing an EC50 of 81.18644 μg / mL, which is the extract of the concentration required to achieve 50% antioxidant activity (Figure 2).

**Figure 2:**
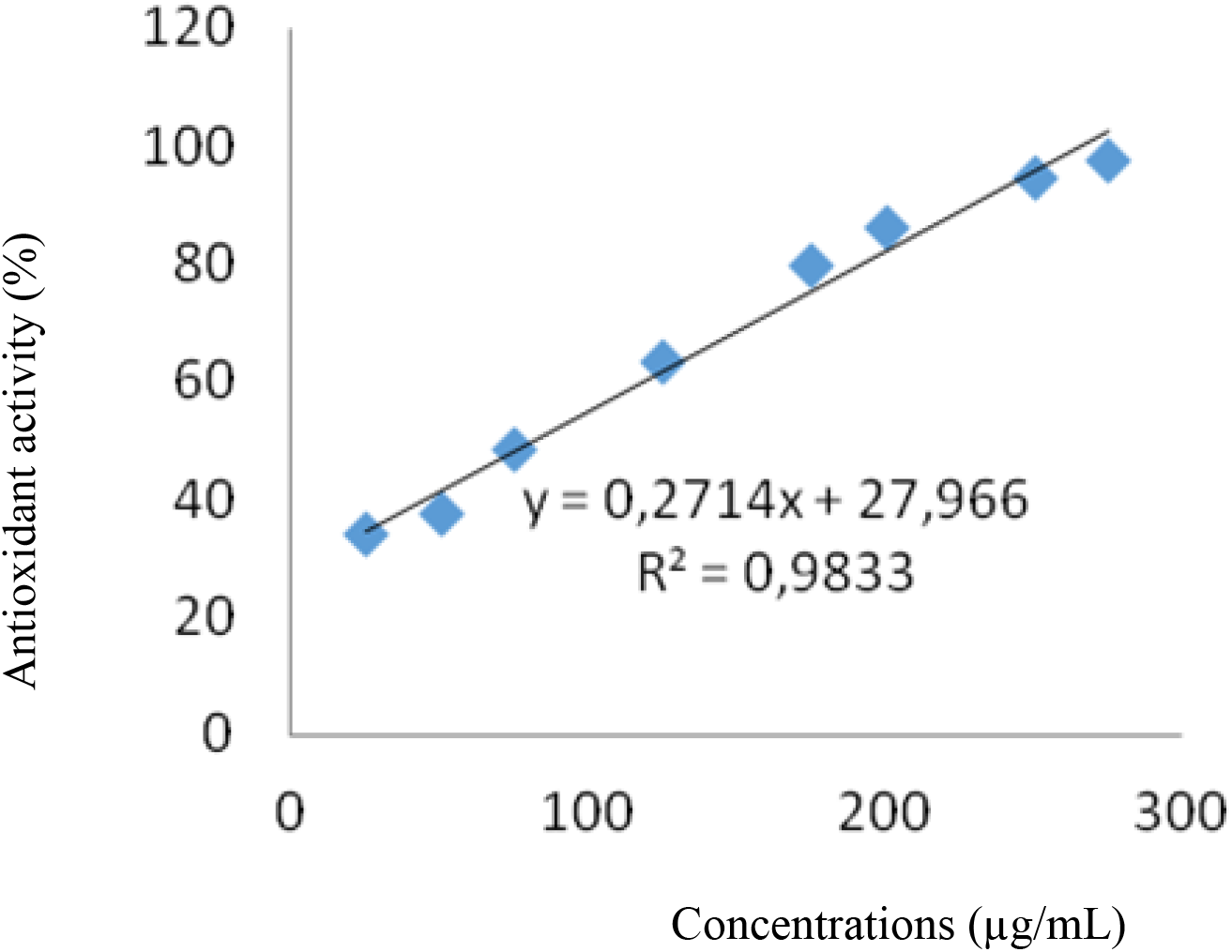
Curve of the percentage of antioxidant activity of the ethanol extract of propolis by the DPPH method.

The total phenolic compound contents calculated by the regression equation y = 0,006x + 0,006, (R2 = 0,999), obtained by the tannic acid calibration curve (where y is the absorbance at 760 nm and x is the concentration of tannic acid in μg / mL) shows that propolis extract has total phenolic contents of 135.33 mg EAT / g (Figure 3).

**Figure 3.**
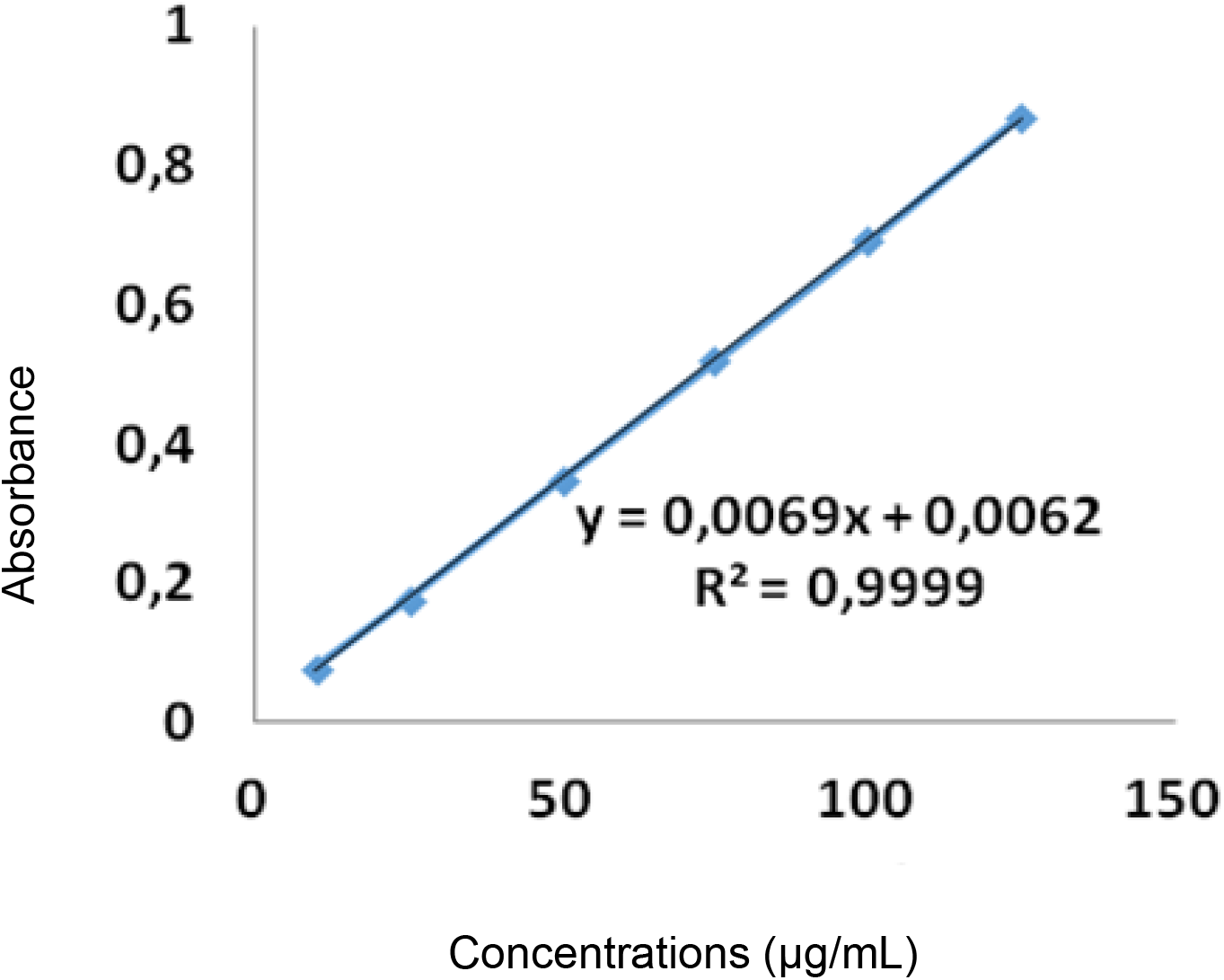
Standard curve of tannic acid for the quantification of phenolic compounds from green propolis extract.

### Antifungal activity of green propolis extract (EPV) against *C. albicans*, *C. parapsilosis* and *C. tropicalis*

The green propolis ethanolic extract inhibited growth of the three *Candida* species evaluated (Table 5). The inhibition halo values of the green propolis ethanolic extract against three *Candida* species are shown in table 5. *C. albicans* and *C. tropicalis* were sensitive to the extracts at 2.5 to 250 μg/mL and *C. parapsilosis* was resistant at the concentrations of 0.25 and 2.5 μg/mL, while at concentrations 25 and 250 μg/mL it was sensitive.

**Table 5.** Antifungal activity of green propolis extract against *Candida* species by disk-diffusion.

| Tested species | Means of the size of the halos (mm)/CIM (µg/mL) of green propolis extract |  |  |  | Control (AFB 16 µg/mL) |
| --- | --- | --- | --- | --- | --- |
|  | 0.25 µg/mL | 2.5 µg/mL | 25 µg/mL | 250 µg/mL |  |
| *C. albicans* | 5 | 15,2 | 17,3 | 20,1 | 25 |
| *C. tropicalis* | 9 | 13,1 | 14,7 | 16,6 | 25 |
| *C. parapsilosis* | 1 | 6,2 | 10 | 12,1 | 10 |

### Adhesion capacity and biofilm formation of *C. albicans*, *C. tropicalis* and *C. parapsilosis* in orthodontic material (acrylic resin and steel)

All *Candida* species were able to adhere and form biofilm on the surfaces of the dental materials studied. In steel and resin, the adhesion intensity of the yeast cells was weak at all incubation times, except for *C. albicans* in 6 and 12h and for *C.parapsilosis* and *C. tropicalis* which presented moderate intensity at 12hs. Regarding biofilm formation (24 and 48 hours), it was observed in steel that *C. albicans* had moderate intensity at 24 and 48 hours; *C. parapsilosis* at 24 and 48 hours had very strong intensity; *C. tropicalis* at 24 hours had strong intensity and at 48 hours very strong. While in the resin, all species at 24 and 48 hours had strong intensity, except for *C. tropicalis* which at 48 hours had very strong intensity (Table 6).

**Table 6.** Adhesion capacity and biofilm formation of *C. albicans*, *C. parapsilosis* and *C. tropicalis* on the surfaces of steel and acrylic resin of orthodontic material according to intensity.

| Time (h) | *Candida* species | Materials |  |  |  |
| --- | --- | --- | --- | --- | --- |
|  |  | Steel |  | Resin |  |
|  |  | Number of adherent cells | Intensity | Number of adherent cells | Intensity |
| 3 | *C. albicans* | 351 | Weak | 161 | Weak |
|  | *C. parapsilopsis* | 175 | Weak | 178 | Weak |
|  | *C. tropicalis* | 236 | Weak | 236 | Weak |
| 6 | *C. albicans* | 693 | Moderate | 580 | Moderate |
|  | *C. parapsilopsis* | 208 | Weak | 209 | Weak |
|  | *C. tropicalis* | 262 | Weak | 331 | Weak |
| 12 | *C. albicans* | 1566 | Strong | 765 | Moderate |
|  | *C. parapsilopsis* | 459 | Weak | 530 | Moderate |
|  | *C. tropicalis* | 610 | Moderate | 520 | Moderate |
|  |  | Number of colonies | Intensity | Number of colonies | Intensity |
| 24 | *C. albicans* | 331 | Moderate | 523 | Strong |
|  | *C. parapsilopsis* | 2435 | Very strong | 554,3 | Strong |
|  | *C. tropicalis* | 913,6 | Strong | 945,6 | Strong |
| 48 | *C. albicans* | 349,3 | Moderate | 578 | Strong |
|  | *C. parapsilopsis* | 1012,3 | Very strong | 920 | Strong |
|  | *C. tropicalis* | 1012,6 | Very strong | 2042,3 | Very strong |

After treatment with ethanolic extract of green propolis, adherence activity of all *Candida* species was reduced compared with the control (saline), showing the efficient activity of green propolis against virulence factors of *Candida* (Figure 4)

**Figure 4.**
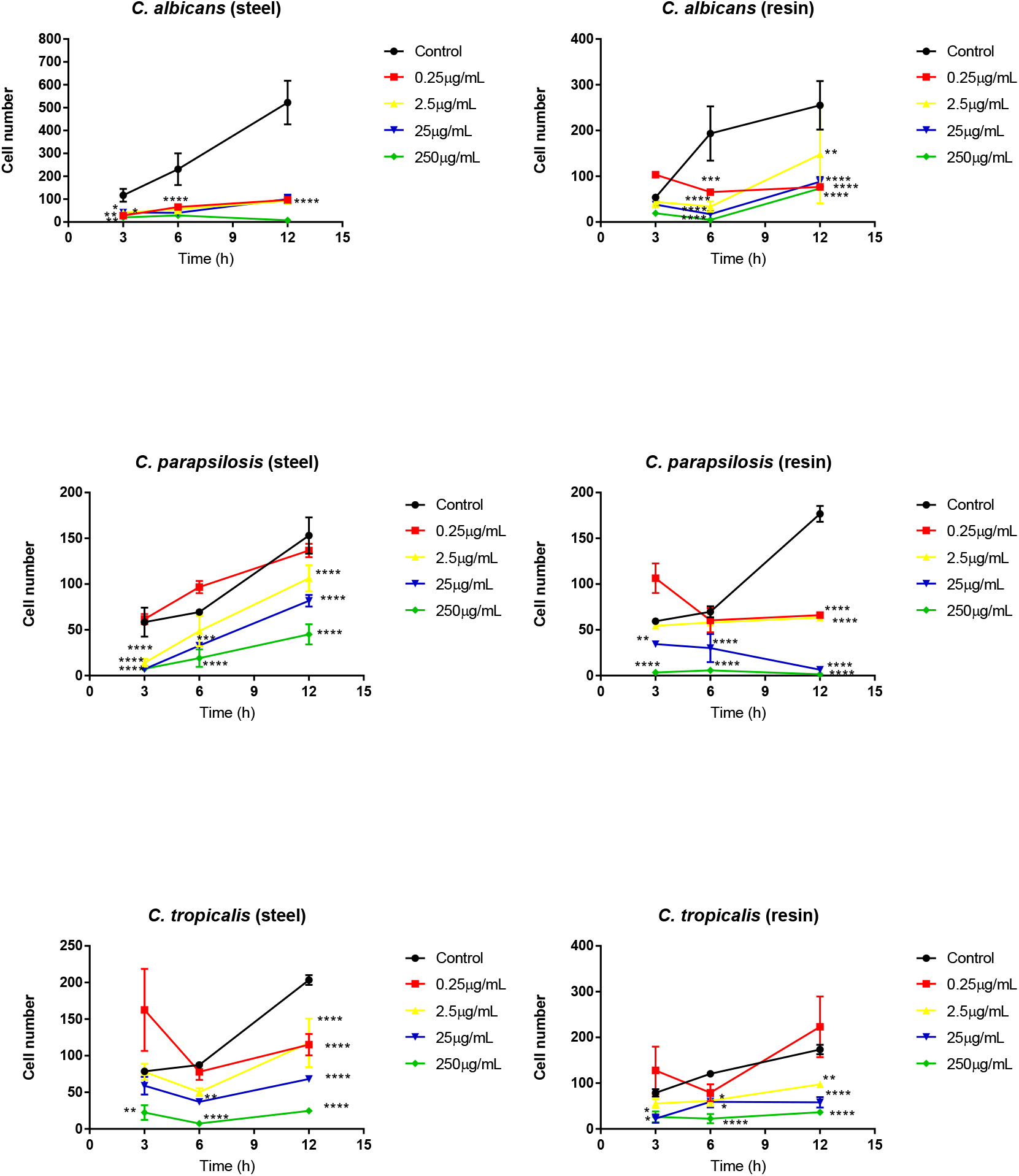
Influence of green propolis extract on the adhesion of *C. albicans*, *C. parapsilosis* and *C. tropicalis* to the surfaces of dental materials (acrylic resin and steel). Effect of extract against *Candida* sp. according to time and material. *p<0.05; **p<0.01;***p<0.001;*p<0.0001

Figure 5 shows the antibiofilm capacity of green propolis. All concentrations have shown the antibiofilm capacity of green propolis extract in 24 and 48 hours.

**Figure 5.**
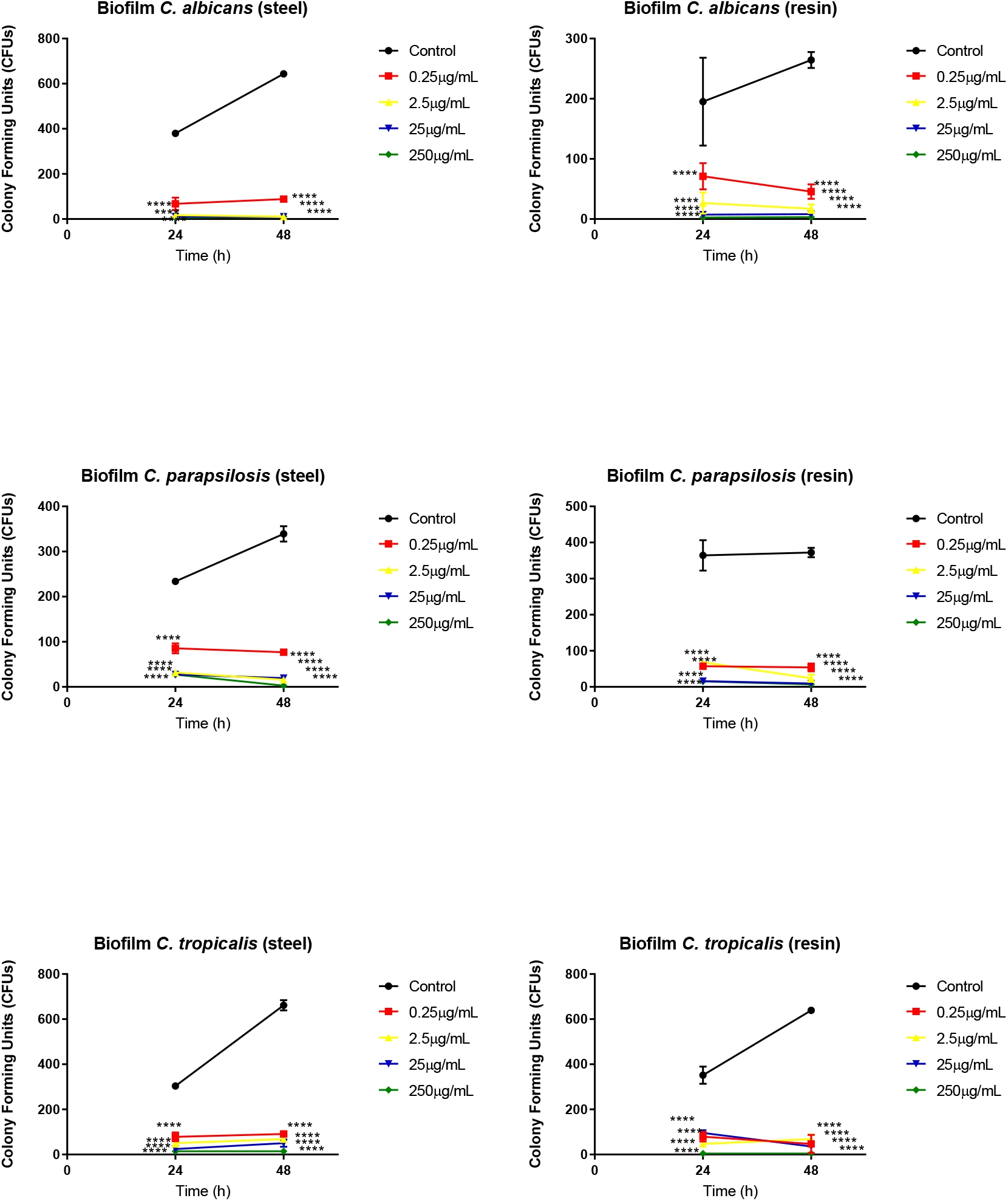
Influence of green propolis extract on the biofilm formation of *C. albicans*, *C. parapsilosis* and *C. tropicalis* to the surfaces of dental materials (acrylic resin and steel). Effect of extract against *Candida* sp. according to time and material. *p<0.05; **p<0.01;***p<0.001;*p<0.0001

## DISCUSSION

The presence of flavonoids, as well as phenolic, aromatic compounds and diterpene acids, in the composition of propolis are associated with several biological properties, such as antifungal (10,13).

The MICs (Minimum Inhibitory Concentrations) of green propolis extract used in the present study against *C. albicans*, *C. parapsilosis* and *C. tropicalis* were 2.5μg / mL. Siqueira et al. (2015) (14) reported 32-64 μg/mL of MIC for red propolis extract, showing the antifungal potential of green propolis extract against this yeast due to its sensitivity to the natural product in a concentration much lower than that found by these authors.

Sforcin et al. (2001) (15) reported that *C. albicans* was more sensitive to propolis from São Paulo (Brazil), located in southeastern Brazil, than *C. tropicalis*. Similar results were found in this study, where *C. albicans* was also more sensitive to propolis than C. tropicalis (the inhibition zones formed by *C. albicans* were larger than those formed by *C. tropicalis*). Propolis antifungal activity against *C. albicans* was studied by Parcker (2007) (16), and D’Auria et al, (2007) (17) where it was suggested that propolis extract inhibits phospholipase extracellular activity, impairing the adhesion of fungal cells to epithelial cells, which was corroborated in the present study (18).

Similar to the results found in this research, propolis extract also showed antibiofilm activity against clinical isolates and ATCC of *Fusarium* species found in patients with onychomycosis, where it was found that the biomass of the treatments decreased significantly when compared to the control, as well as the number. of viable cells (19).

Capoci et al. (2015) (20) observed a reduction of more than 50% of CFUs for all *Candida albicans* isolates after exposure to Propolis Extract (PES) compared to control. These results corroborate those found in this research, where there was also a reduction of CFUs of *C. albicans, C. tropicalis* and *C. parapsilosis* at 25 and 250 μg / mL at all abiotic materials tested.

In this study, a greater reduction in *C. albicans* biofilm formation was observed at all concentrations compared to the biofilms produced by *C. parapsilosis* and *C. tropicalis*, and the reduction of biofilm in *C. parapsilosis* was significantly higher from the concentration of *C. parapsilosis*. 25 μg/mL and *C. tropicalis* at 250 μg/mL, corroborating the work of Tobaldini-Valerio et al. (21) who also observed a greater biofilm reduction (~ 3.5 log) in *C. albicans*, followed by *C. parapsilosis* and *C. tropicalis*, with a reduction of approximately 2.8 and 2 log, respectively, at all Propolis Extract concentrations. tested.

The Green Propolis Ethanol Extract (EEPV) used in this study showed fungicidal, anti-adherent and antibiofilm activity on *C. albicans*, *C. parapsilosis* and *C. tropicalis* on dental materials (steel and acrylic resin) at the concentration of 2.5 μg/mL, suggesting the preventive use of this natural product in oral infections by the genus *Candida*.

## MATERIALS AND METHODS

### Obtaining and Preparation of the Green Propolis Ethanolic Extract (EEPV)

The green propolis used in the *in vitro* assays was acquired from the Rosita Apiary (Betim-MG). Fresh propolis was stored in a dry, airless plastic bag kept under refrigeration until use. The hydroalcoholic extract of green propolis was obtained according to the methodology of Soares de Moura et al. (2011) (22). Approximately 200g of green propolis was diluted in 500 ml of PA ethyl alcohol, stored in an amber flask and stored at room temperature with stirring for 2h/day for 8 days. It was then filtered and rotaevaporated at 35 ° C until complete solvent removal. The resulting concentrate was lyophilized and stored refrigerated until use.

### Phytochemical screening

The extract was submitted to phytochemical screening based on the methodology presented by Matos (2009) (13) to detect phenols and tannins (reaction with ferric chloride); anthocyanins, anthocyanidins, flavonoids, leucoanthocyanidins, catechins and flavanones (pH variation using hydrochloric acid and sodium hydroxide); flavonols, flavanones, flavanonols and xanthones (reaction with metallic magnesium and concentrated hydrochloric acid). The results obtained in each test were qualitatively evaluated by staining reactions and precipitate formation.

### Determination of total phenolic compounds

The determination of total phenolics of the extract occurred by the Folin-Ciocalteu method based on procedures described by Waterhouse (2012) (23), with some modifications.

### Gallic acid standard curve

For the determination of the standard curve of tannic acid, a solution of 2,000 μg.mL^−1^ was prepared which gave five different dilutions (10, 25, 50, 75, 100, 125 μg tannic acid mL-1). Thereafter, 500μL of each solution was diluted with 2.5 mL of 10% (v / v) Folin-Ciocalteu solution and 2 mL of 4% (m / v) sodium carbonate solution, then mixed in test tubes. This mixture was protected from light. After 30 minutes, the absorbance was read on a spectrophotometer at 760 nm using a quartz cuvette. The absorbance readings were plotted as a function of gallic acid concentration through the regression equation and its coefficients.

### Evaluation of antioxidant activity by DPPH (2,2-Diphenyl-1-picrylhydrazy)

The antioxidant activity of the extracts was evaluated with 1,1-diphenyl-2-picrilidrazil (DPPH), according to the methodology described by Yen and Wu (1999) (24). From the extract concentrations (10, 25, 50, 75, 100, 125, 150, 175, 200 and 225 μg / mL) a reaction mixture with DPPH was prepared. Subsequently, 1.0 mL of each dilution was transferred to a test tube containing 3.0 mL of DPPH ethanolic solution (0.004%). After 30 minutes of incubation in the dark at room temperature, DPPH free radical reduction was measured by reading the absorbance using a 517 nm spectrophotometer. A blank sample was prepared using ethanol instead of the sample. Equation 1 was used to calculate the ability to sequester the free radical expressed as a percentage of radical oxidation inhibition.

Antioxidant activity (%) = [1- (Sample Absorbance / Control Absorbance] x 100.

The IC50 value (concentration of the extract needed to sequester 50% of DPPH radical) was calculated by the above equation based on the concentrations of the extracts and in their respective percentages of DPPH radical sequestration. The analyzes were performed at the Chemical Research Laboratory of the Federal University of Maranhão.

### Analysis of the phytochemical composition

The analysis of the phytochemical composition of the extract was obtained by High Performance Liquid Chromatography (HPLC) coupled to mass spectrometer (HPLC-DAD MS). Chromatographic analyzes were performed at the Instrumentation Analytical Center of the Institute of Chemistry of the University of São Paulo. After solubilization, the hydroalcoholic extract of green propolis was analyzed by high performance liquid chromatography (HPLC). Shimadzu^®^ chromatograph (Shimadzu Corp. Kyoto, Japan) consisting of a solvent injection module with a Shimadzu LC-20AD pump and Shimadzu UV-Vis detector (SPDA-20A) was used for analysis. The column used was Supelco Ascentis C-18 (250 x 4.6 mm - 5um). HPLC was performed with an elution gradient using a mobile phase with water and 5% acetic acid and methanol (organic phase) in different proportions. The total time of the experiment was 115 minutes. The injection volume was 20 μL and chromatographic acquisition was performed at 270 nm (DAD). Data were collected and processed using LC Solution software (Shimadzu). Identification of compounds by mass spectrometry was performed in negative mode.

### Dental Material and Microorganisms

Fragments of dental material from Self-Curing Acrylic (Resin) and Orthodontic Band (Metal) types were purchased from dental shops. Three species of *Candida* were used in this study: *Candida albicans* ATCC 443-805-2, *Candida parapsilosis* ATCC 726-42-6 and *Candida tropicalis* ATCC 1036-09-2 obtained from the stock collection of the Collection of Fungi of Immunology and Mycology Laboratory - NIBA/UFMA.

### Evaluation of EEPV antifungal activity

Initially *Candida* species were cultivated on Sabouraud Agar incubated at 37°C in a BOD greenhouse and after 24 hours were diluted in saline according to McFarland at a 05. scale. Antifungal activity was performed by the disc diffusion method on Muller-Hinton agar with 2% dextrose and 0.5 μg / mL methylene blue as recommended by the CLSI M44-A2 protocol (2009) (25). Amphotericin B was diluted in PBS 1x plus 1% DMSO to give a concentration of 16 μg / mL for positive control. To evaluate antifungal activity, 50mg of EPP was diluted in 500μl of DMSO, from this dilution it was prepared a working solution by diluting 1ml in 9ml of PBS 1x, from which concentrations of the extract of 0.25, 2.5, 25 and 250μg / mL were obtained. Inhibition halos were evaluated according to interpretation criteria of CLSI (2009) (25) and Capocci (2013) (20).

**Table 7.**
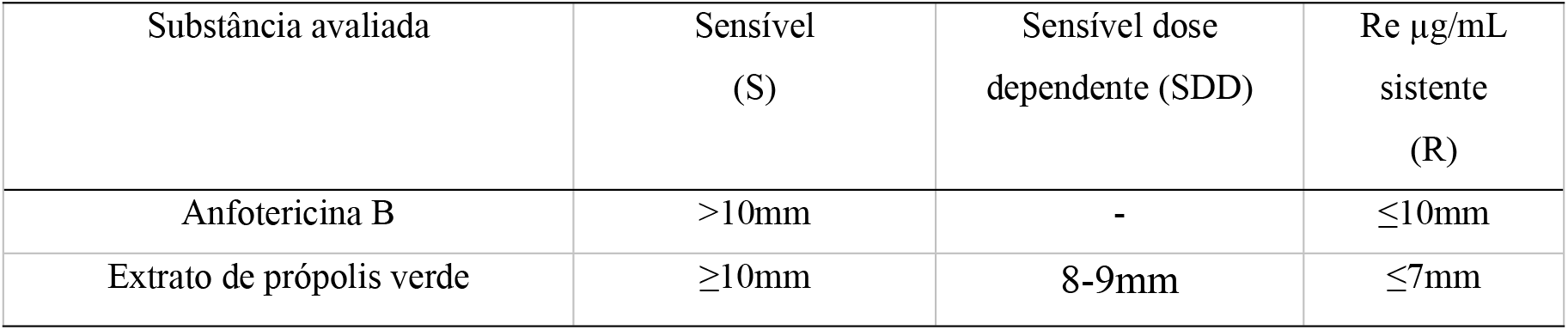
Interpretation criteria of fungi susceptibility to green propolis extract and amphotericin B by disk-difussion (CLSI M44-A2; CAPOCCI, 2013).

| Substância avaliada | Sensível (S) | Sensível dose dependente (SDD) | Re µg/mL sistente (R) |
| --- | --- | --- | --- |
| Anfotericina B | >10mm | - | ≤10mm |
| Extrato de própolis verde | ≥10mm | 8-9mm | ≤7mm |

### Adherence and biofilm formation on abiotic metal and acrylic resin surfaces

5 cm-sized fragments of dental material (metal and acrylic resin) were made and used in this study as described by Silva et al (2010) (26) and Borges et al. (2018) (27) with modifications. The fragments were cultivated in saline with 100μl of a 1×10^4^ cel/mL suspension of *C. albicans*, *C. parapsilosis* and *C. tropicalis* and they were kept in a BOD greenhouse at times of 3, 6 and 12hs for adherence and 24 and 48hs for biofilm in triplicate. Then the fragments were washed 3x with sterile distilled water, fixed with PA alcohol and stained with violet crystal. Subsequently the fragments were added to tubes containing 3 ml 0.85% saline and vortexed for 10 minutes obtaining a fungal suspension of the cells adhered to the materials. 10μl of the adherence test suspension was added in a Neubauer chamber to count the adherent cells under light microscopy and according to the number of cells quantified the intensity of adhesion to the dental material was classified as: Negative: <50 yeast / ml; Weak: between 50 and 499 c / ml; Moderate: 500 to 999 c / ml; Strong: 1000 or more c / ml. For the biofilm test, 100μl of the suspension was added to a plate containing Muller-Hinton Agar to quantify the number of colony forming units (CFUs) and the biofilm intensity formed in the dental material was classified as: Negative: without CFU growth; Weak: growth between one and 199 CFUs, Moderate: from 200 to 499, Strong: with 500 to 1000 CFUs and Very strong: over 1000 CFUs.

### Anti-adherent and antibiofilm activities of EEPV

EEP dilutions (0.25, 2.5, 25 and 250μg / mL) were prepared as described above. To evaluate the effect of the extract, the fragments were cultivated in a tube containing 3 ml of each concentration of EEP and incubated in a BOD greenhouse at 37°C at times of 3, 6 12hs for adhesion and 24 and 48hs for biofilm. After each period, the tubes were removed from the greenhouse, the fragments were washed with 3X sterile distilled water, and after each wash and greenhouse drying, the fragments were fixed with PA ethyl alcohol and stained with violet crystal. Then the fragments were added in saline tube and vortexed for 10 minutes.

### Statistical analysis

The data were analyzed using the program “GraphPad Prism R” version 7. A two-way ANOVA with Tukey post-hoc was performed, where p <0.05 and a confidence interval of 95% were considered.

## Acknowledgments

The authors would like to acknowledge FAPEMA for providing financial support. For Post-graduate Program in Adult Health from Federal University of Maranhão.

## Supplementary Data

### Chemical compounds identified by HPLC-DAD-MS

**Fig S1.**
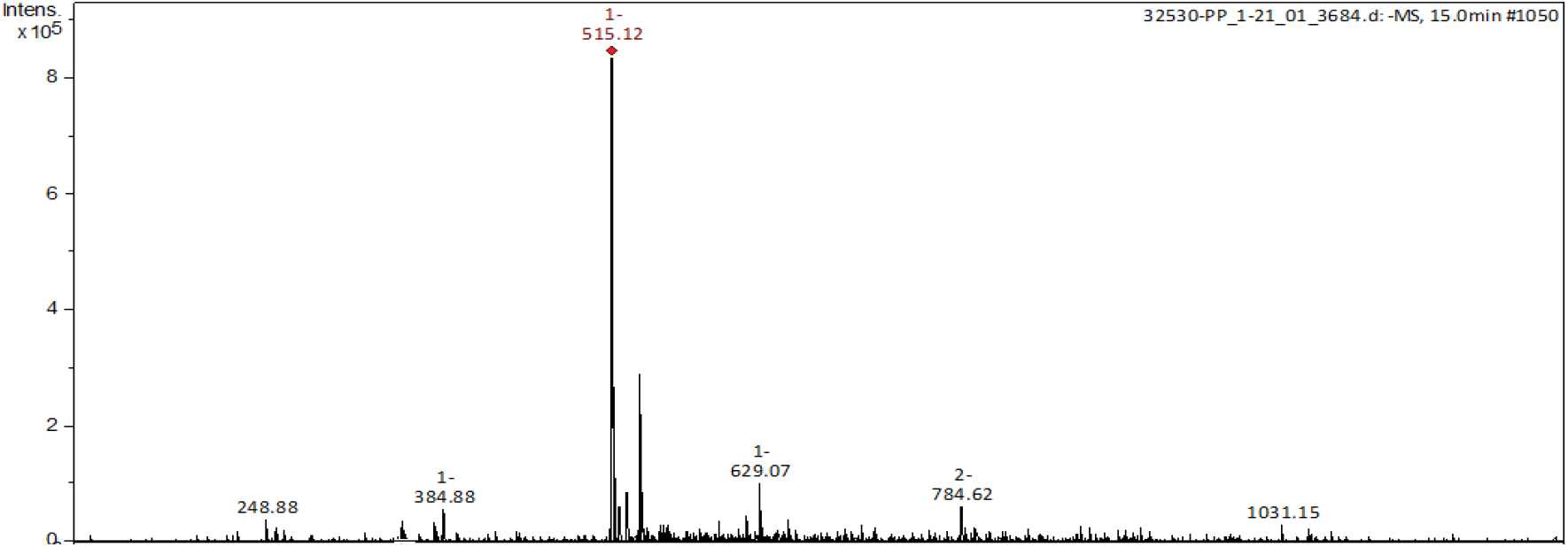
Mass Spectrum of [M–H]^−^ for 1

**Fig S2.**
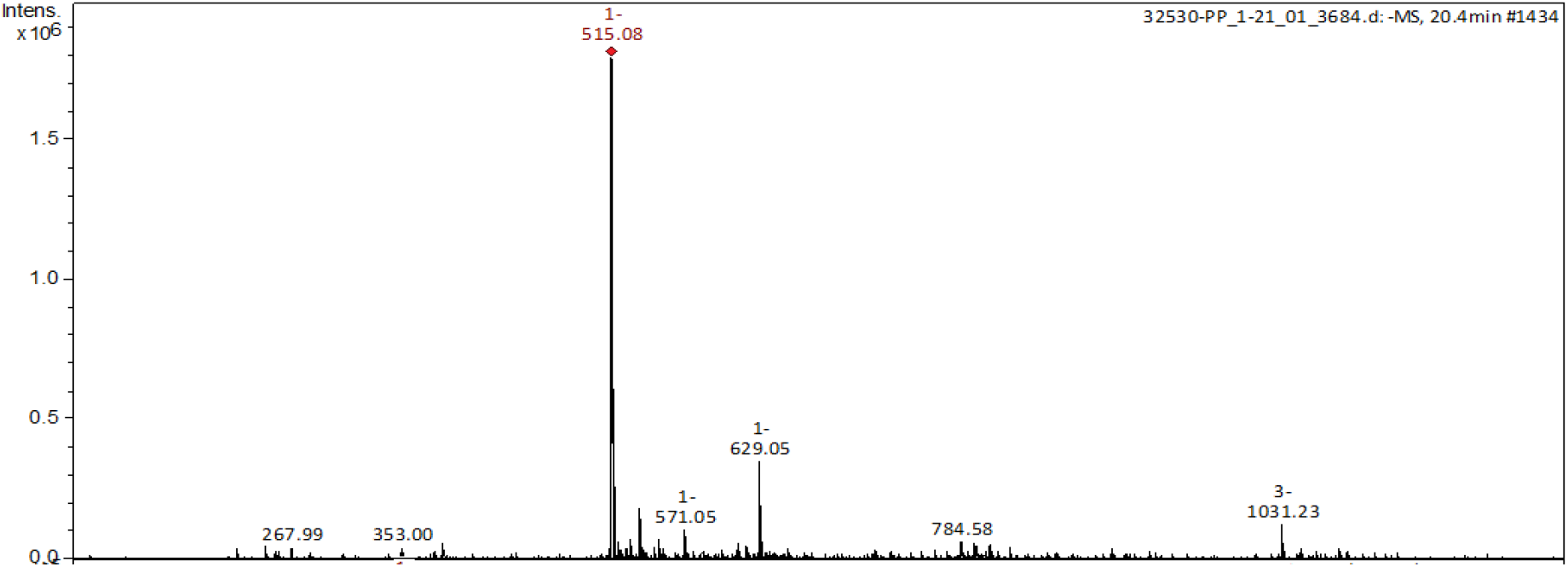
Mass Spectrum of [M–H]^−^ for 2

**Fig S3.**
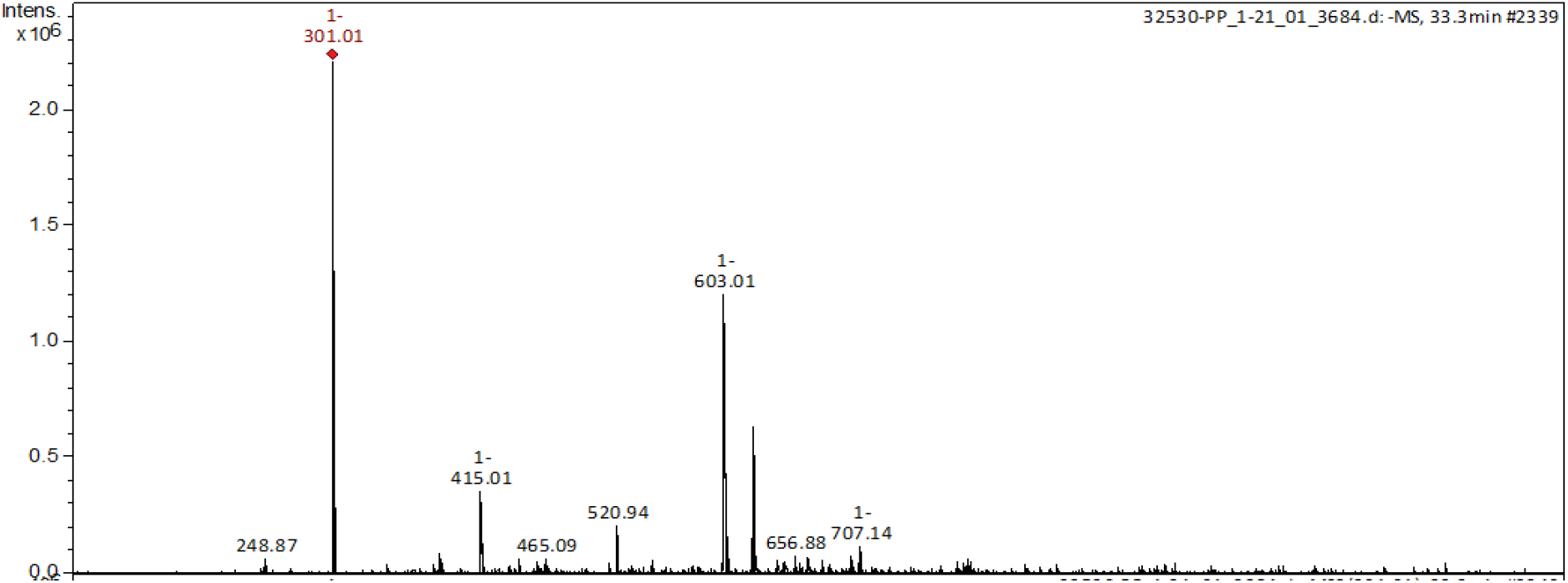
Mass Spectrum of [M–H]^−^ for 3

**Fig S4.**
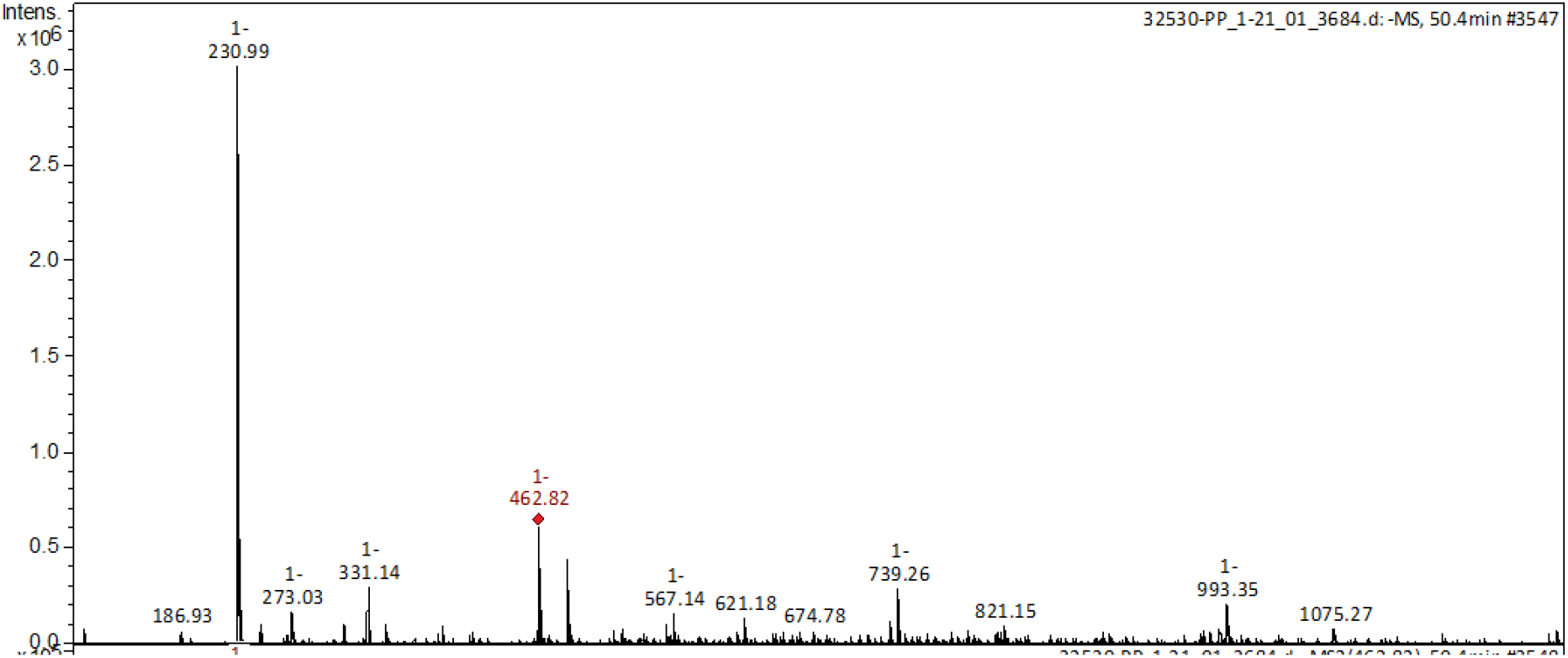
Mass Spectrum of [M–H]^−^ for 4

**Fig S5.**
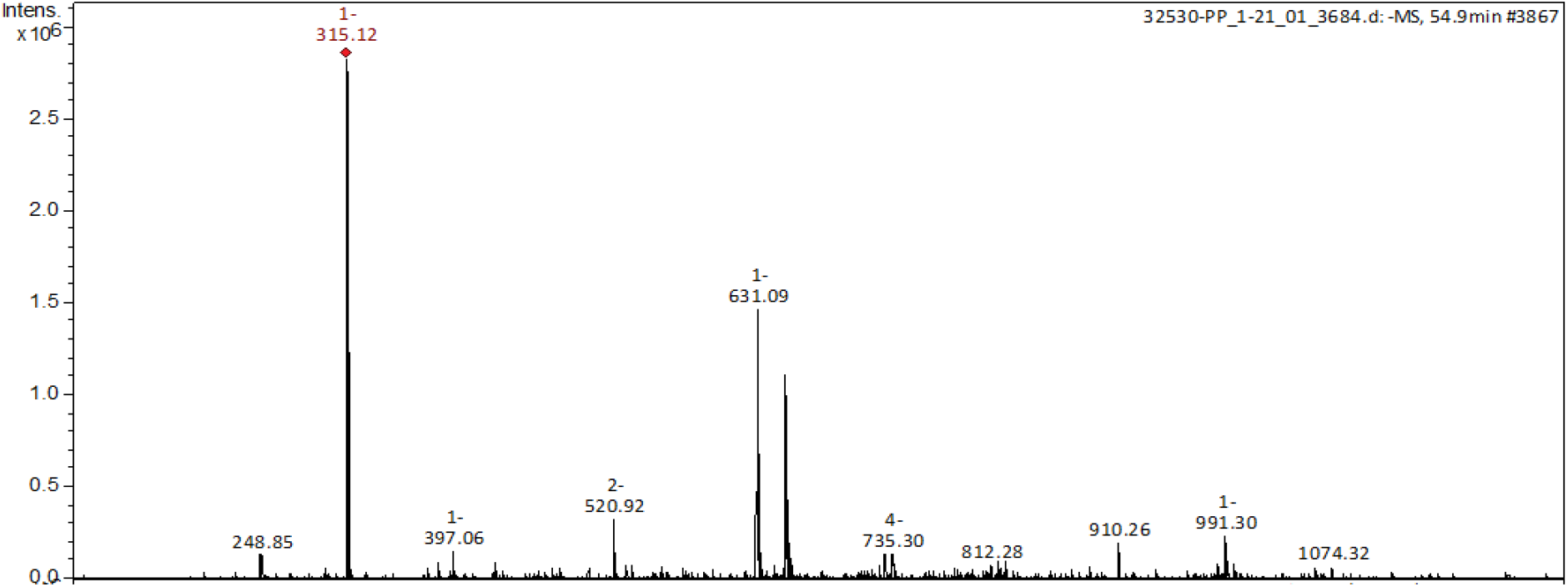
Mass Spectrum of [M–H]^−^ for 5

**Fig S6.**
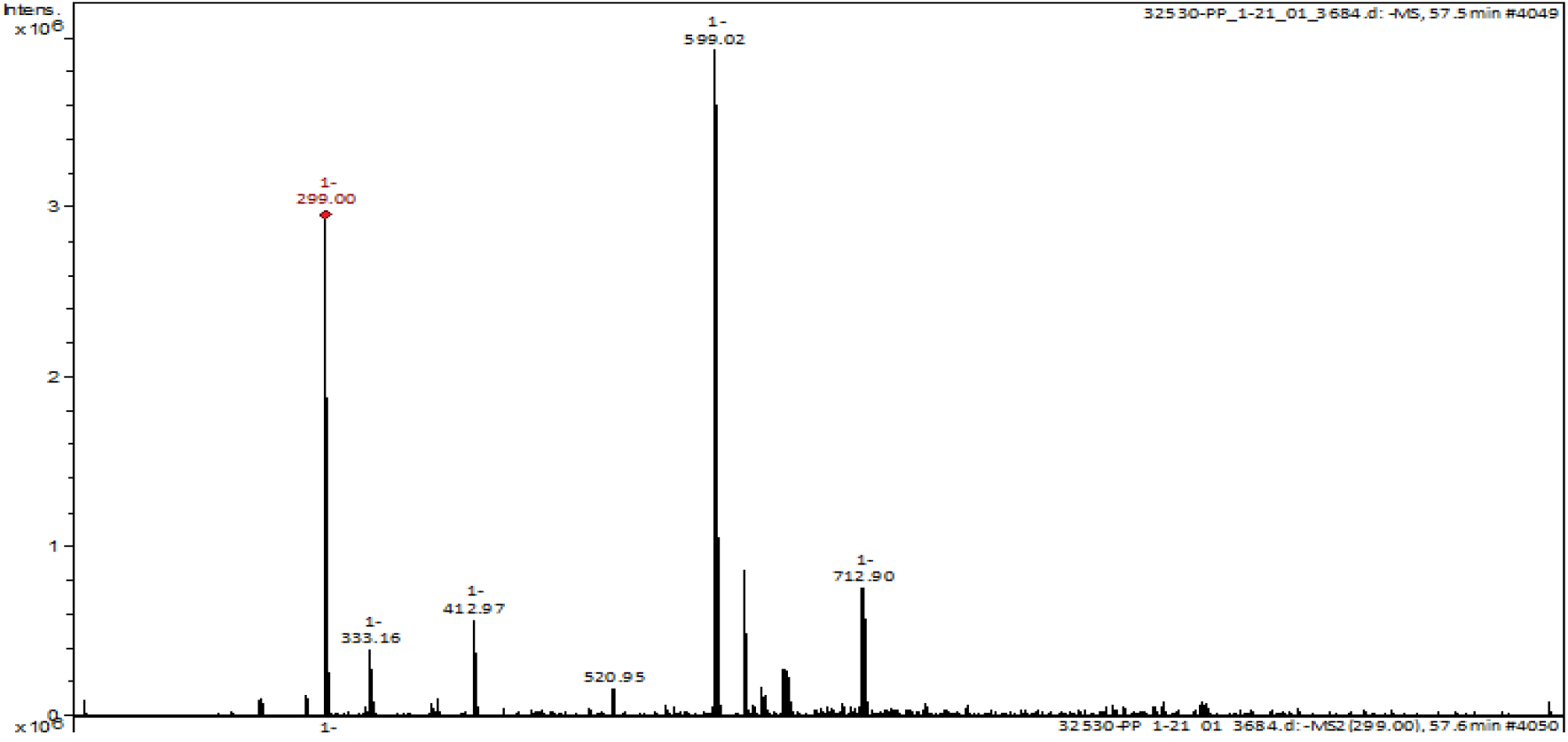
Mass Spectrum of [M–H]^−^ for 6

**Fig S7.**
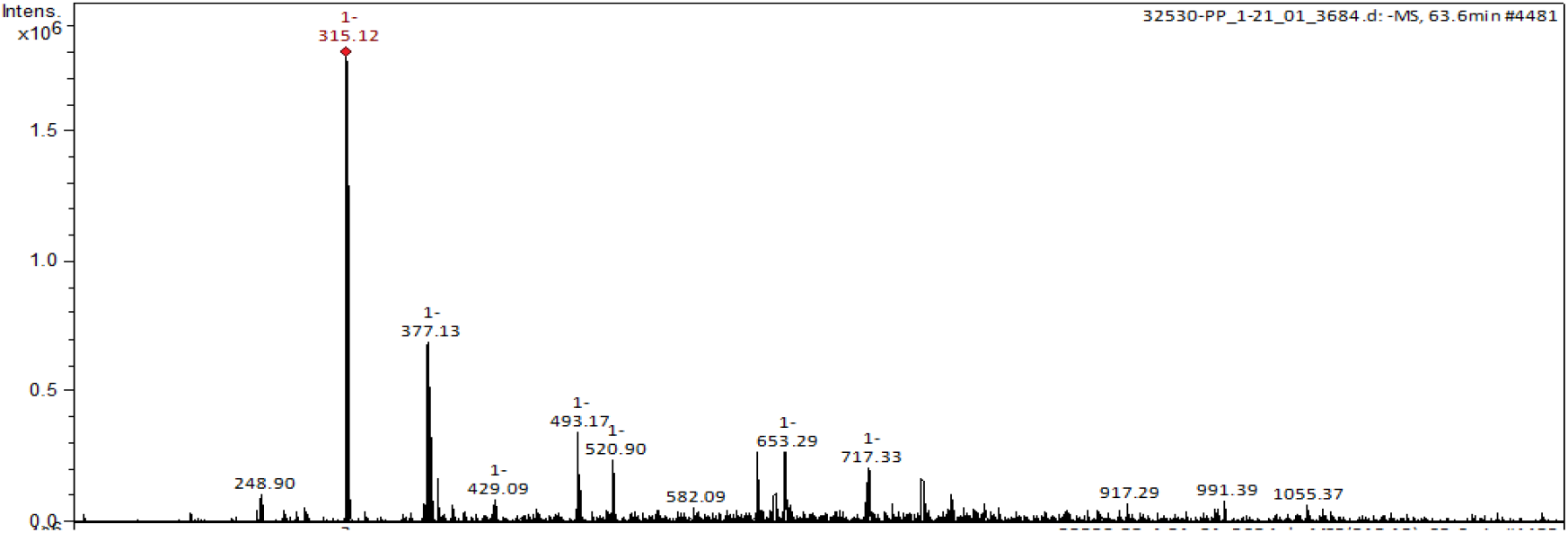
Mass Spectrum of [M–H]^−^ for 7

**Fig S8.**
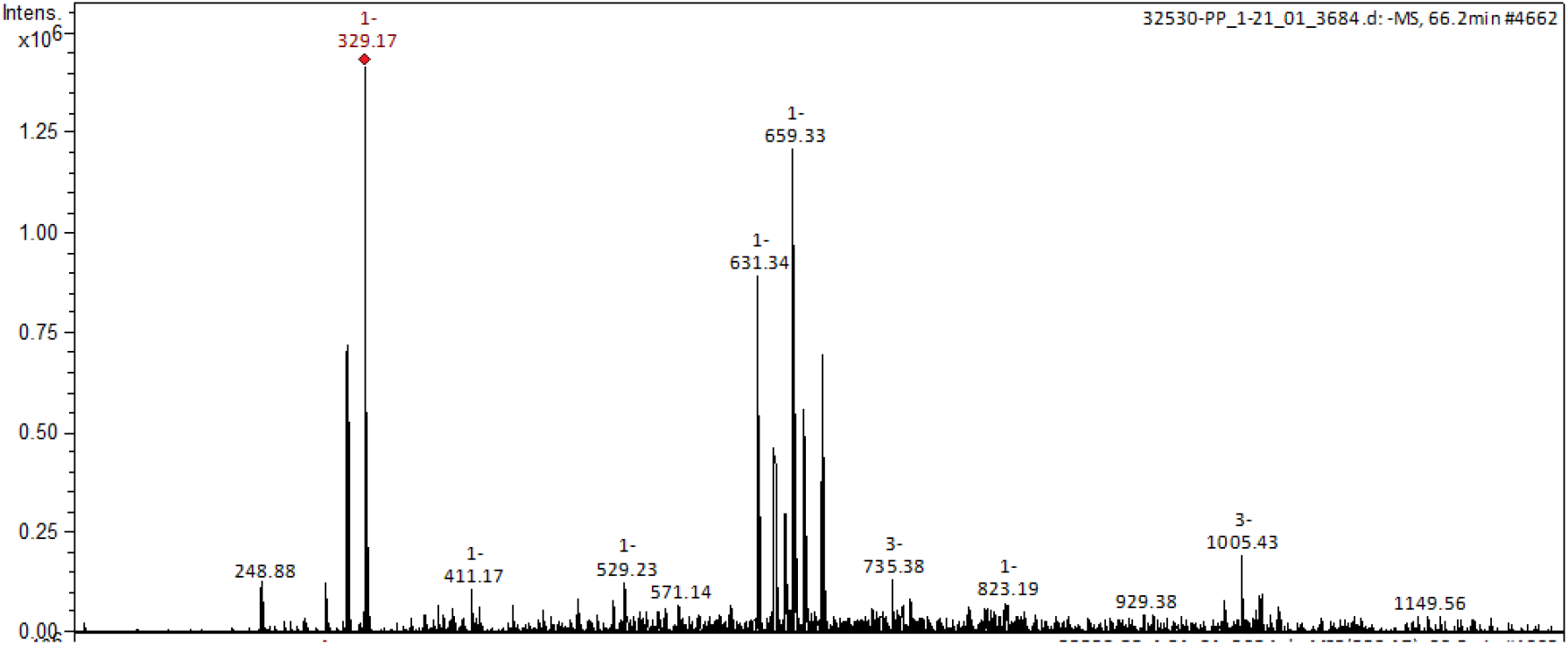
Mass Spectrum of [M–H]^−^ for 7

**Fig S9.**
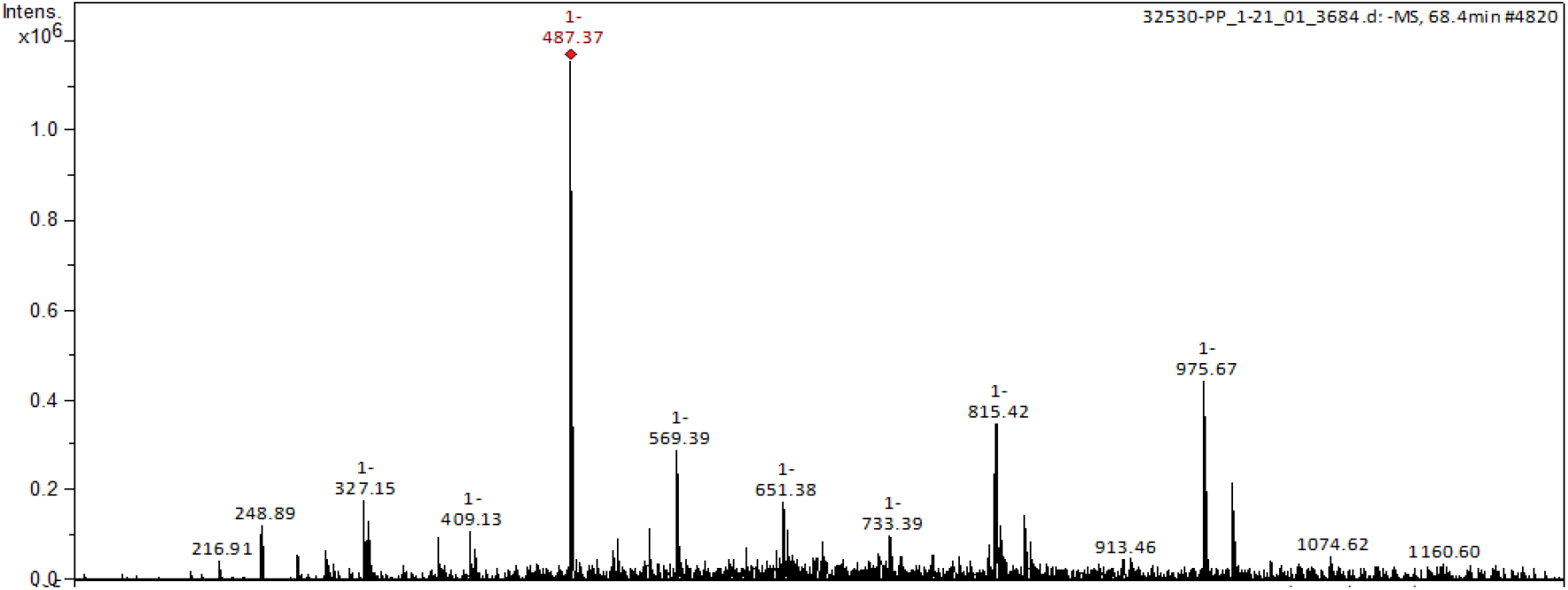
Mass Spectrum of [M–H]^−^ for 8

**Fig S10.**
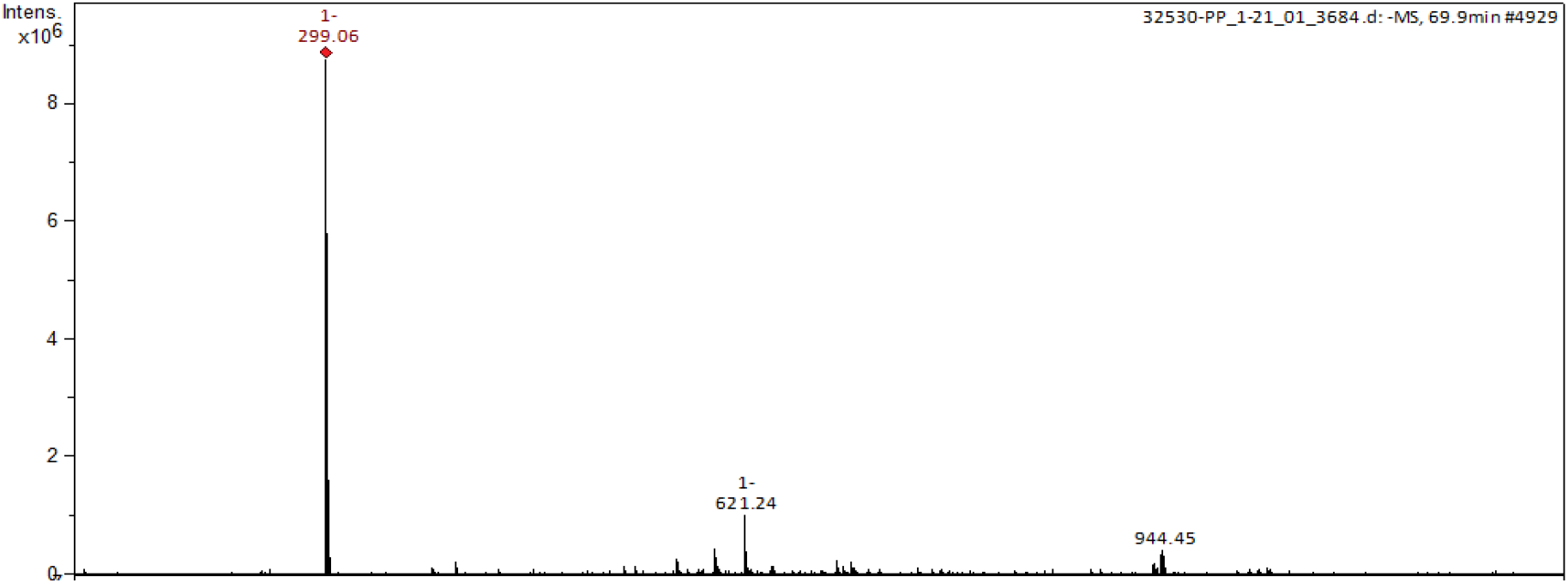
Mass Spectrum of [M–H]^−^ for 10

**Fig S11.**
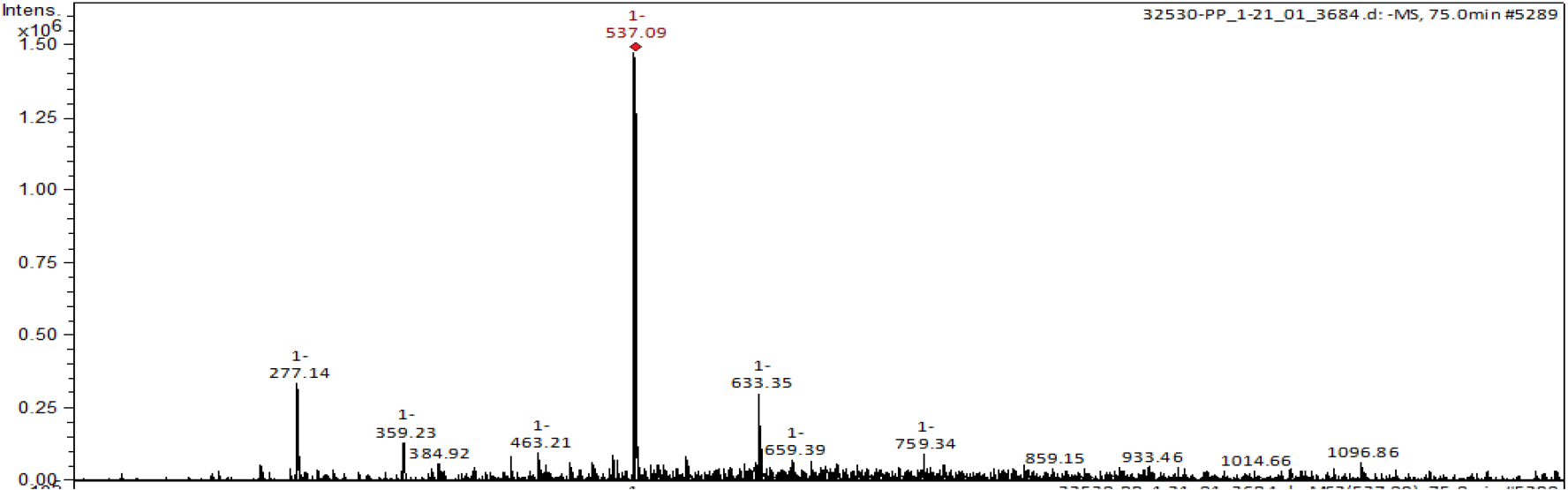
Mass Spectrum of [M–H]^−^ for 11

**Fig S12.**
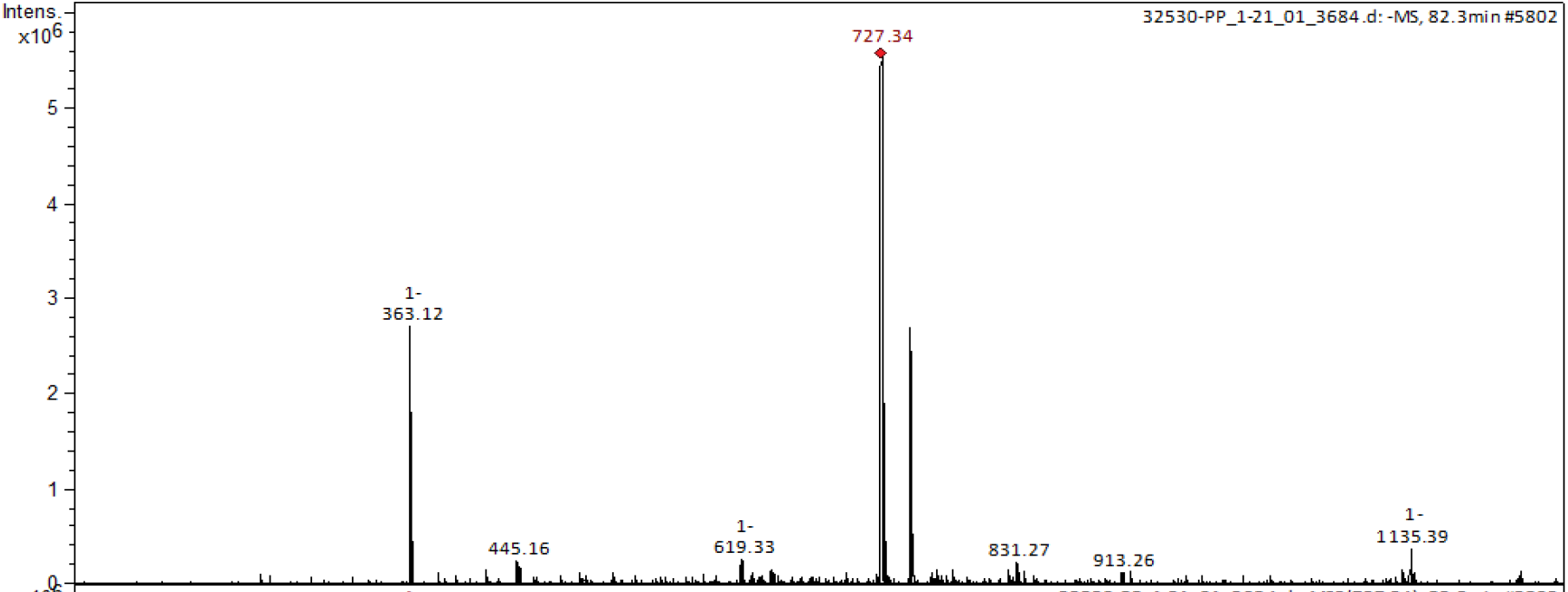
Mass Spectrum of [M–H]^−^ for 12

**Fig S13.**
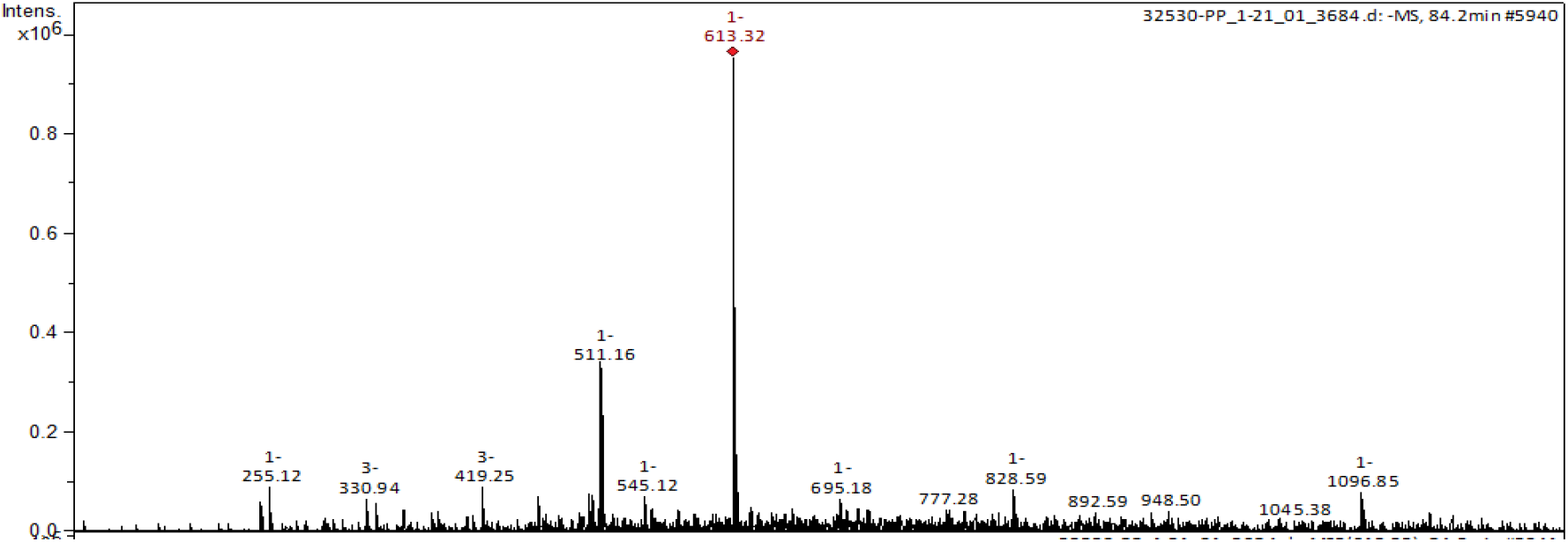
Mass Spectrum of [M–H]^−^ for 13

**Fig S14.**
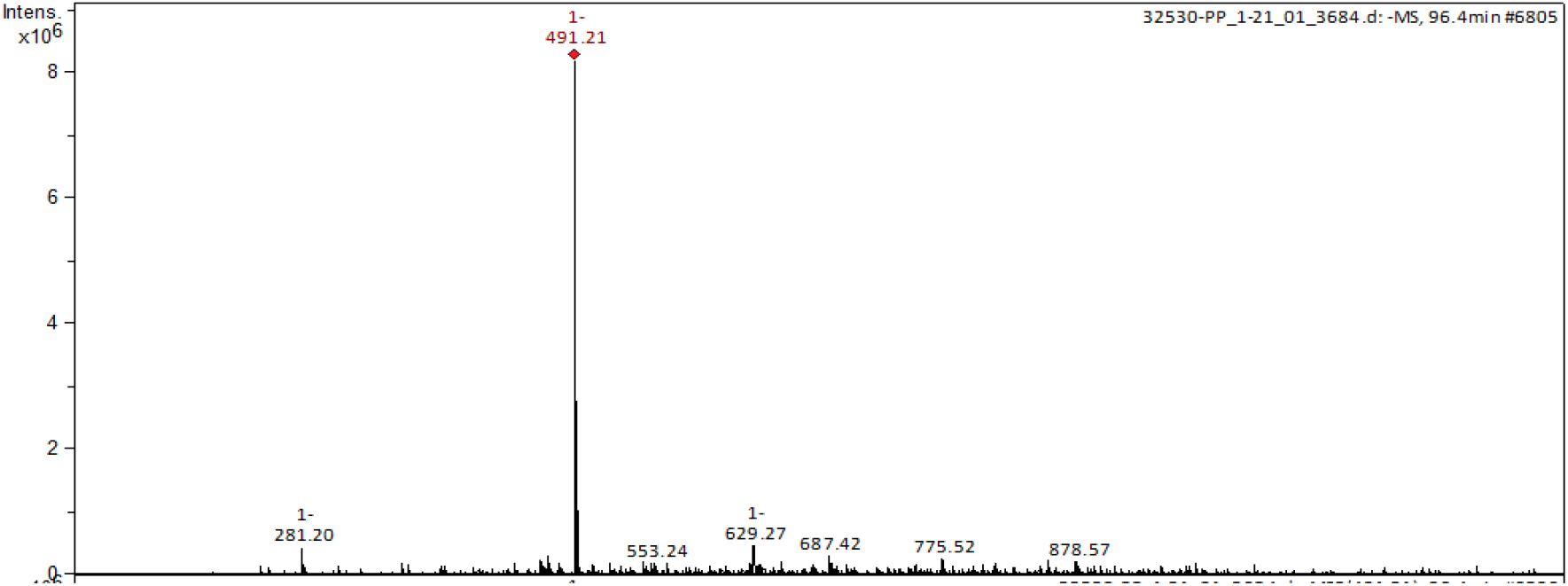
Mass Spectrum of [M–H]^−^ for 14

**Fig S15.**
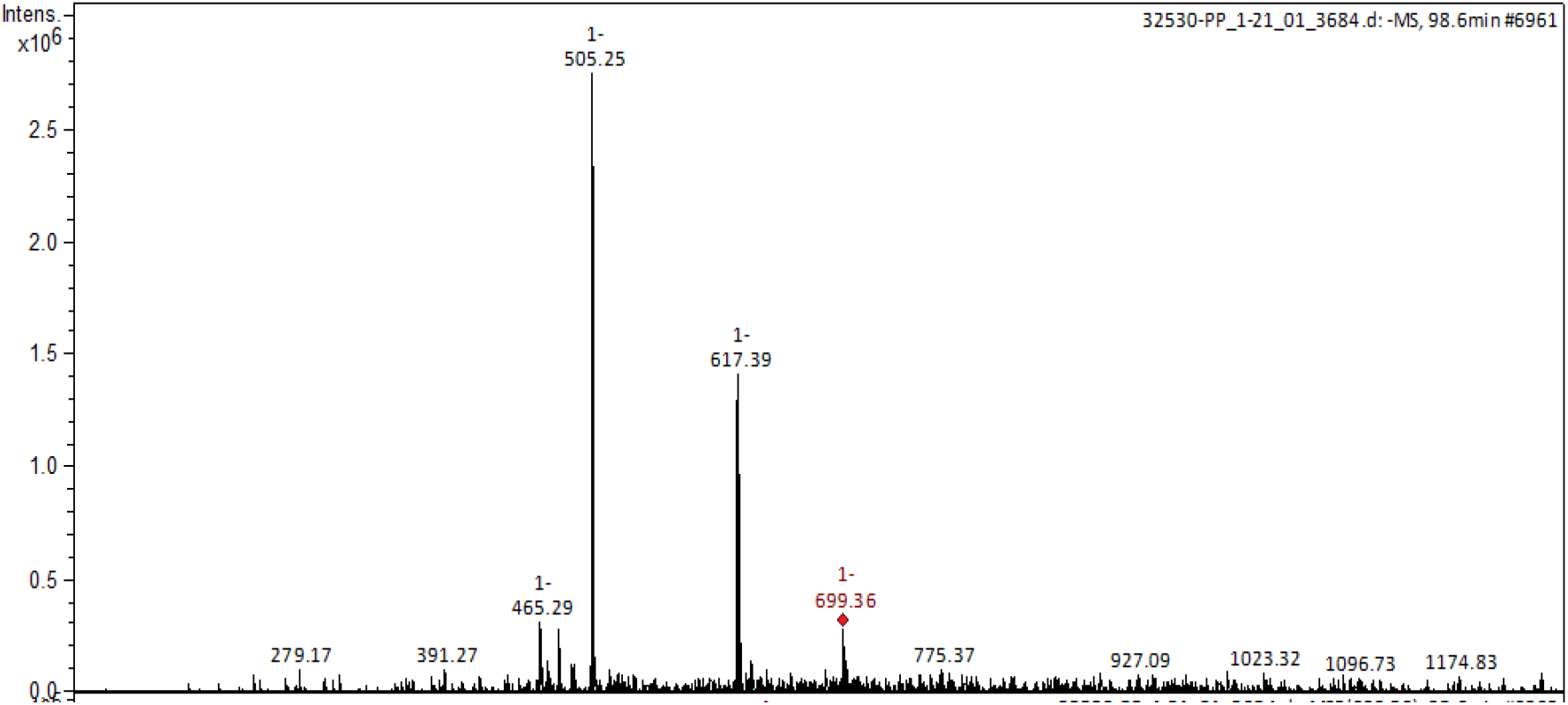
Mass Spectrum of [M–H]^−^ for 15

